# Use of Next Generation Sequencing to study two cowpox virus outbreaks

**DOI:** 10.1101/444141

**Authors:** Markus H. Antwerpen, Enrico Georgi, Alexandra Nikolic, Gudrun Zöller, Peter Wohlsein, Wolfgang Baumgärtner, Christophe Peyrefitte, Remi Charell, Hermann Meyer

## Abstract

**Background:** Between 2008 and 2011 about 40 cases of human cowpox were reported from Germany and France. Infections had been acquired via close contact to infected, young pet rats. Sequencing of the hemagglutinin gene of various cowpox virus (CPXV) isolates resulted in an identical and unique sequence in each case pointing to a common source. In a second CPXV outbreak in cats in a small animal clinic in Germany in 2015, four out of five hospitalized cats showed identical hemagglutinin sequences and thus, a hospital-acquired transmission was assumed.

**Methods:** Homogenates of lesion material from rats, cats and humans were cultivated in cell culture. The genomes of 4 virus isolates, 9 CPXVs from our strain collections and from DNA of 3 paraffin-embedded lesion materials were determined by Next Generation Sequencing (NGS). For phylogenetic analyses a MAFFT-alignment was generated. A distance matrix based on concatenated SNPs was calculated and plotted as dendrogram using Unweighted Pair Group Method with Arithmetic mean (UPGMA) for visualization.

**Results:** Aligning of about 200.000 nucleotides of 8 virus isolates associated with the pet rat outbreak revealed complete identity of six genomes, the remainder two genomes differed in as little as 3 SNPs. When comparing this dataset with four already published CPXV genomes also associated with the pet rat outbreak, again a maximum difference of 3 SNPs was found. The outbreak which lasted from 2008 till 2011 was indeed caused by a single strain which has maintained an extremely high level of clonality over 4 years.

Aligning genomic sequences from 4 cases of feline cowpox revealed 3 identical sequences and one sequence which differed in 65 nucleotides. Although identical hemagglutinin sequences had been obtained from four hospitalized cats, genomic sequencing proved that a hospital-acquired transmission had occurred in only three cats.

**Discussion:** Analyzing the rather short sequence of the hemagglutinin gene is not sufficient to conduct molecular trace back analyses. Instead, whole genome sequencing is the method of choice which can even be applied to paraffin-embedded specimens.

**Funding Statement:** This publication was supported by the European Virus Archive goes Global (EVAg) project that has received funding from the European Union’s Horizon 2020 research and innovation program under grant agreement No 653316.

This study was also supported in part by the European Union’s Horizon 2020 research and innovation program under grant agreement No 643476 (COMPARE).

The funders had no role in study design, data collection and analysis, decision to publish, or preparation of the manuscript.

## Introduction

The genus *Orthopoxvirus*, family *Poxviridae*, subfamily *Chordopoxvirinae*, presently contains four species which are pathogenic to humans: variola virus (VARV), the causative agent of smallpox, monkeypox virus (MPXV), vaccinia virus (VACV), and cowpox virus (CPXV). Mousepox (ECTV) and camelpox virus (CMLV) can affect mice and camels, respectively. Including tatera poxvirus (TATV), these species are collectively referred to as “Old World” orthopoxviruses (OPVs). In addition, three species of North American origin, i. e. raccoon, vole and skunk poxvirus, are currently known. [1].

While naturally occurring VARV infections are indeed no longer of concern – except in a bioterrorist context ‐ human infections with MPXV and CPXV are increasingly reported. While cases of human monkeypox are restricted to the tropical rainforest of Africa, CPXV is endemic in Western Eurasia. Although initially seen in cows, it is now generally accepted that the species CPXV is misnamed [2]. Rodents, particularly vole species, are regarded to be the true reservoir hosts from which spillover infections to accidental hosts are regularly observed. Accidental hosts are cows, domestic cats, horses and also exotic animals in zoos, such as large felidae and elephants, which regularly develop severe disease. Humans in direct contact with infected accidental hosts, especially cats and pet rats, can develop local lesions, in rare cases a generalization is possible [3].

Historically, OPVs have been differentiated from other poxviruses by demonstration of a hemagglutinin (HA)-positive phenotype able to agglutinate chicken erythrocytes. Within the genus *Orthopoxvirus* single species had been differentiated based on biological characteristics, including mouse virulence, size and color of pocks on the chorioallantoic membrane as well as formation of A-type inclusion bodies. A pathognomonic feature of isolates assigned to the species CPXV is the induction of hemorrhagic pocks on the chorioallantoic membrane and the formation of intracytoplasmatic A-type inclusion bodies [4] However, conducting these assays is laborious and time consuming. Therefore, attempts had been made to correlate the phenotype with the genotype of OPV species either by generation of Restriction Fragment Length Polymorphism data or by sequencing single genes, such as the hemagglutinin (HA) gene[5]. Both genome-based methods successfully confirmed the current species concept originally based on phenotypic characteristics. Sequencing of the HA gene had become the method of choice to assign a given OPV isolate to a species of the genus *Orthopoxvirus* and numerous HA sequences from various cases are available in GenBank [3], [6]–[12]. As compared to VARV and MPXV, the HA sequences of CPXVs are much more diverse. This genetic variation can be explained by the existence of local variants which have evolved over time in a rodent reservoir [2]. Recently, results of phylogenetic analyses of 58 full-length CPXV genomes confirm the polyphyletic character of the species CPXV which consists of at least four different clades and several variants [6], [7]. The classification of CPXV strains into clades roughly followed their geographic origin. A rather large outbreak occurred in Germany and France during 2008 and 2011 with about 40 cases of human cowpox virus infection [10], [11], [14]. All humans had acquired their infection via young pet rats. Most of the pet rats had died rather quickly due to pulmonary infection caused by CPXV. Characterization of 14 viruses/clinical materials by sequencing the respective HA genes proved a 100% identity [8], [10], [11], [14] indicating that a single CPXV strain circulating in rat breeder facilities could have been the common source. Recently an atypical manifestation of CPXV infection in 5 cats in the larger area of Hannover was described by Jungwirth [15]. Identical HA gene sequences were identified in 4 cats, and in combination with epidemiological data a hospital-acquired transmission was assumed [15].

With the introduction of benchtop sequencers, enabling even small routine diagnostic laboratories to perform whole genome sequencing, sequencing of only a part of the microorganisms’ genome is not a riddle anymore. Determination of drug resistances or virulence genes can be performed easily based on the same sequence dataset, as well as identification and *in-silico* typing and clonal assignment of the investigated strains.

In this study Next Generation Sequencing (NGS) was used to re-analyze the pet rat and the feline cowpox outbreak. We determined the genomes of further strains and lesion materials and conclude that the pet rat-associated outbreak which lasted from 2008 till 2011 was indeed caused by a unique CPXV strain. In addition, we determined genomic sequences of the 2015 Hannover outbreak in cats and conclude that hospital acquired cowpox infections had occurred in three out of four hospitalized cats.

## Methods

### Viruses

Four clinical specimens suspected to contain CPXV were received in Germany in 2009. Two specimens were collected from a 13-year old boy and his pet rat; the boy presented with local skin lesions and his rat died with severe pulmonary symptoms. One specimen was lesion material from a boa which had been bitten by a feeder rat, which had been bought in a pet store in Marl, Germany. The fourth specimen originated from a rat, which had bought in the same pet store in Marl and the rat had died showing severe respiratory symptoms. Homogenates of the lesions were prepared in cell culture medium using a bead-beater and aliquots were inoculated onto African Green Monkey Kidney cell line MA 104 as described recently [3]. Four viral strains, each corresponding to a specific specimen, were isolated and designated as strains “Boy Biederstein”, “Rat Biederstein”, “Rat Marl” and “Boa Marl”, respectively (Table 1). CPXV identification was achieved through sequencing the PCR product corresponding to the entire ORF of the HA gene [14].

**Table 1:**
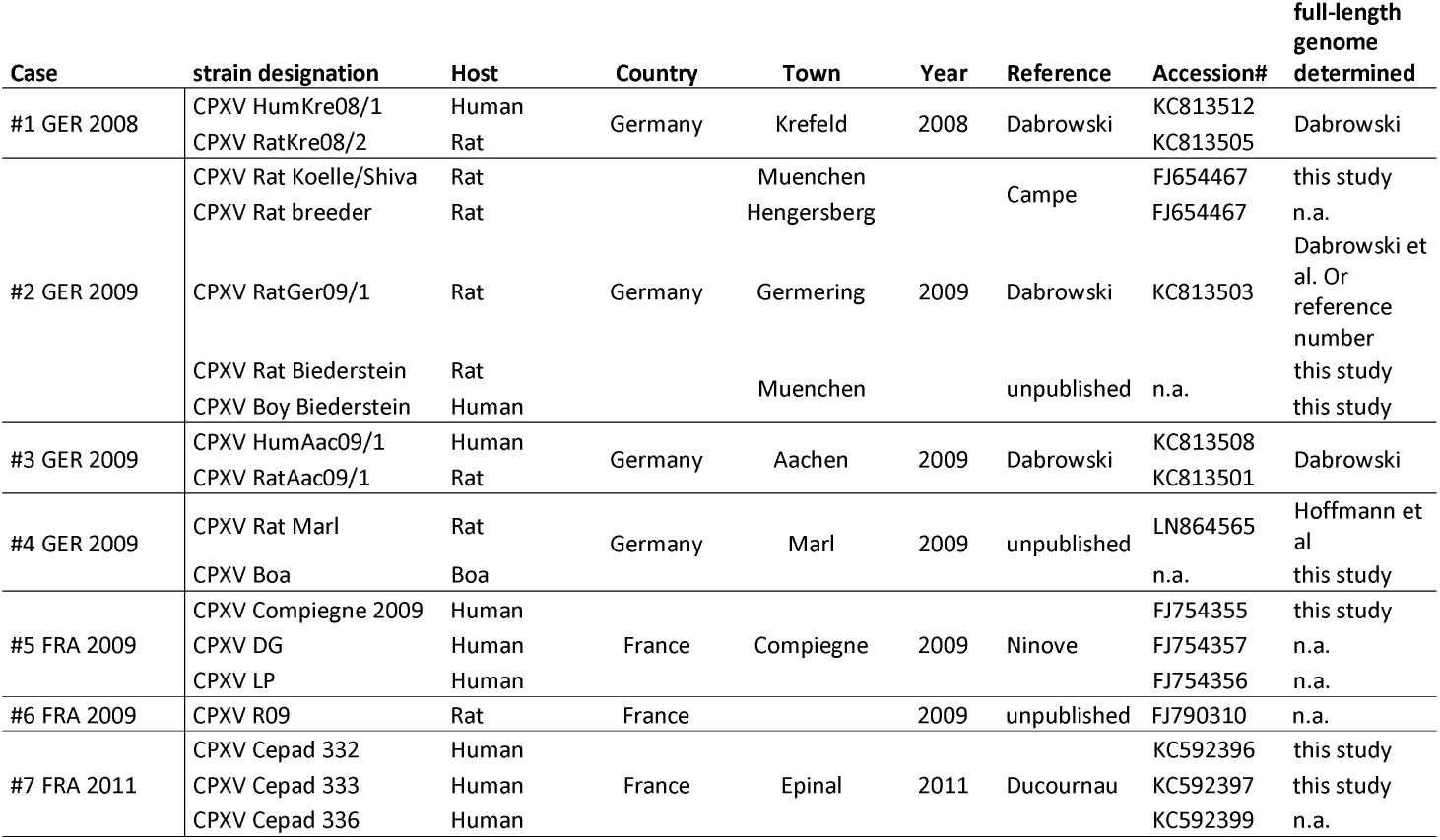
List of cases associated with the pet rat outbreak 2008 till 2011 (HA-sequence group C030; Table _S1) (See last pages)

| Case | strain designation | Host | Country | Town | Year | Reference | Accession# | full-length genome determined |
| --- | --- | --- | --- | --- | --- | --- | --- | --- |
| #1 GER 2008 | CPXV HumKre08/1<br>CPXV RatKre08/2 | Human<br>Rat | Germany | Krefeld | 2008 | Dabrowski | KC813512<br>KC813505 | Dabrowski |
| #2 GER 2009 | CPXV Rat Koelle/Shiva | Rat | Germany | Muenchen | 2009 | Campe | FJ654467 | this study |
|  | CPXV Rat breeder | Rat |  | Hengersberg |  |  | FJ654467 | n.a. |
|  | CPXV RatGer09/1 | Rat |  | Germering |  | Dabrowski | KC813503 | Dabrowski et al. Or reference number |
|  | CPXV Rat Biederstein<br>CPXV Boy Biederstein | Rat<br>Human |  | Muenchen |  | unpublished | n.a. | this study<br>this study |
| #3 GER 2009 | CPXV HumAac09/1<br>CPXV RatAac09/1 | Human<br>Rat | Germany | Aachen | 2009 | Dabrowski | KC813508<br>KC813501 | Dabrowski |
| #4 GER 2009 | CPXV Rat Marl | Rat | Germany | Marl | 2009 | unpublished | LN864565 | Hoffmann et al |
|  | CPXV Boa | Boa |  |  |  |  | n.a. | this study |
| #5 FRA 2009 | CPXV Compiagne 2009 | Human | France | Compiagne | 2009 | Ninove | FJ754355 | this study |
|  | CPXV DG | Human |  |  |  |  | FJ754357 | n.a. |
|  | CPXV LP | Human |  |  |  |  | FJ754356 | n.a. |
| #6 FRA 2009 | CPXV R09 | Rat | France |  | 2009 | unpublished | FJ790310 | n.a. |
| #7 FRA 2011 | CPXV Cepad 332 | Human | France | Epinal | 2011 | Ducournau | KC592396 | this study |
|  | CPXV Cepad 333 | Human |  |  |  |  | KC592397 | this study |
|  | CPXV Cepad 336 | Human |  |  |  |  | KC592399 | n.a. |

### Whole genome sequencing

Genomic DNA was prepared from virus-infected cell cultures. A total of eight CPXV strains associated with the pet rat outbreak were analyzed. Four strains were part of our strain collections (“Compiegne 2009”, “Rat Koelle”, “Cepad 332”, and “Cepad 333”) and four strains had been isolated (strains “Rat Marl”, “Boa Marl”, “Rat Biederstein” and “Boy Biederstein”; Table 1). With regard to the feline cowpox outbreak [15] a total of five CPXV strains (“75/01”, “Moritz 2015/3a”, “Sammy 2015/4a”, “Leo 2015/5” and “S2216/04”) of our collections were investigated (Table 2).

**Table 2:**
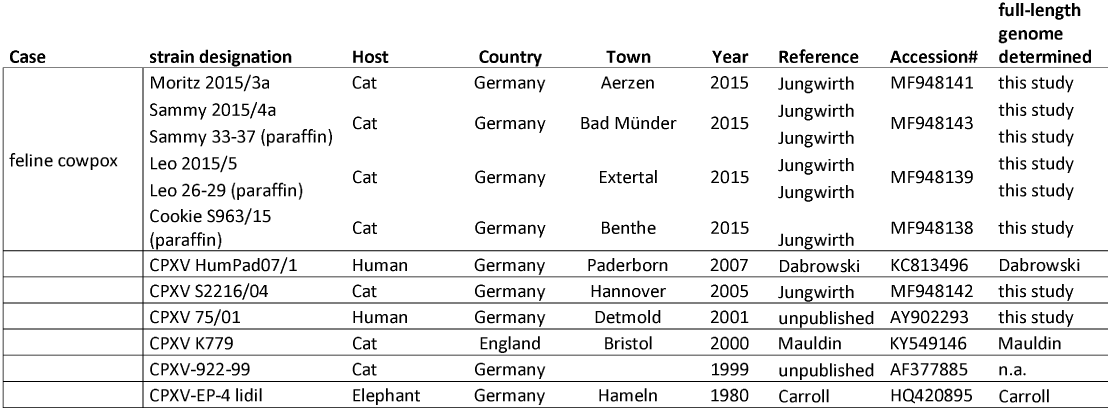
Cases of feline cowpox and strains with identical HA-sequences (HA-sequence group C029; Table _S1)

Infected cell cultures were freeze-thawed three times and genomic DNA was extracted by using the automated DNA extraction system MagNA Pure Compact (Roche Diagnostics, Penzberg, Germany). Genomic DNA was also extracted directly from paraffin-embedded skin tissues of cats Sammy, Leo and Cookie using the FFPE-DNAeasy kit (Qiagen, Hilden, Germany).

DNA quantification was performed using the fluorometric system Qubit 2.0 (Life Technologies, Darmstadt, Germany) and the Qubit^®^ dsDNA high sensitivity assay kit (Thermo Fisher Scientific). Nextera^®^ XT DNA Library Preparation kit (Illumina) with an input DNA amount of 3-5 ng was used for library preparation. Finally, whole genome sequencing was performed on a MiSeq instrument (Illumina) with corresponding MiSeq Reagent Kit v3 (600 cycle) chemistry.

### Bioinformatics

Sequences encoding the entire open reading frame (ORF) of the hemagglutinin gene of “Old World” OPVs were downloaded from NCBI nucleotide archive. The respective strains (n= 462) and their accession numbers are listed (Table S1). For sequence clustering a Unweighted Pair Group Method with Arithmetic mean (UPGMA)-analysis was performed using software Bionumerics and visualized as Minimum-Spanning-tree (MST).

### Sequence Analysis

The sequenced paired-end reads were mapped against chromosomes of *Catus felis* (RefSeq NC_018723 ‐ NC_018741) or *Chlorocebus sabaeus* (RefSeq NC_023642 ‐ NC_02364272) in order to discard sequence reads of eukaryotic background DNA. Remaining reads were assembled de-novo with SPAdes v3.9 [16]. Hence, resulting contigs were aligned against CPXV reference strain Brighton Red (RefSeq NC_003663.2) using blastn [17] to determine their order. Afterwards, an in-house developed Python script was used to concatenate sequences into one single scaffold by resolving overlaps or creating gaps filled with Ns of a length estimated from the mapping. BWA [18] was subsequently used to map the raw reads to the chromosomal contigs. Finally, Pilon [19] was applied to undo mis-assemblies, polish the sequence by correcting SNPs and small InDels, as well as to close gaps in order to obtain one continuous chromosome based on almost the complete whole genome sequence. The Genome Annotation Transfer Utility (GATU [20]) was used for annotation of genomes. The criteria for annotation included 90% nt similarity with CPXV strain Brighton Red. After the transfer of annotation, assigned open reading frames of all genomes were visually inspected and corrected manually, if needed. All generated genomes were deposited into NCBI GenBank as a BioProject <u>PRJNA369073</u>.

For phylogenetic analyses, a MAFFT-alignment [21] of all sequenced and downloaded reference virus genomes (Table 1 and 2) was generated using the FFT-NS-2 algorithm and inspected and manually corrected using MEGA 6.0, if needed. Using an in-house python script, only those positions present in all datasets were used for further calculations. Furthermore, for determining of SNP-calling, only those SNPS were used, of which within a surrounding area of 10 nucleotides no further SNPs were observed in the used dataset, and no gap in the alignment for at least one sequence was observed. A Maximum Parsimony distance matrix using the accelerated transformation (ACCTRAN) was calculated and plotted as dendrogram using R (packages Phangorn and Phytools). Bootstrap values were calculated with 1000 replicates.

## Results

### Hemagglutinin-based phylogenetic relationships of Orthopoxviruses

We reconstructed the phylogeny of Old World OPVs by using sequences covering the entire ORF of the HA gene since the number of HA sequences by far exceeded those of other genes. As of June 2018, a total of 462 sequences were retrieved from Genbank and aligned. Details about the species, the year and location of isolation, the host and the length of the ORFs are given in Table S1. A total of 152 unique sequences were identified and identical sequences were assigned common group numbers. Bayesian inference showed a consensus Minimum-Spanning-Tree (Figure 1). Circle sizes are relative to the number of identical sequences and branch lengths are relative to the number of nucleotide changes. In general, all sequences of strains and isolates belonging to the species MPXV (n=112 sequences), VARV (n=87), CMLV (n=39) and ECTV (n=11) clustered closely together with little variation. The four species are clearly separated from each other, thus reflecting accurately the current taxonomy of the genus *Orthopoxvirus*.

The species VACV is monophyletic as well, however, variation is much higher with as many as 34 different variants out of 52 reported sequences. This can be best explained because VACV vaccine strains have been repeatedly passaged in various cell cultures in different laboratories. Since there was no selective pressure to survive in a reservoir host, *in vitro* replication has favored evolution into pools of highly diverse quasi-species [22]. The genetic variation of VACVs described here mirrors results of recent full-genome analysis [23]. The authors conclude that VACV was once a circulating virus in animals in Europe in the 19th century that later disappeared in this reservoir.

Comparative analyses of CPXV sequences display a much higher genetic diversity: 144 strains can be allocated to 79 unique sequence groups. In contrast to the *orthopoxvirus* species mentioned above, phylogenetic analyses clearly demonstrate that CPXV cannot be regarded as a monophyletic species: there are several different clades as already reported by [6]–[8]. However, some strains might even be erroneously attributed to VACV while indeed belonging to CPXV. The genetic diversity of CPXVs is also reflected by different lengths of the HA ORF, ranging from 816 bp to 963 bp. A recent phylogenetic analysis of 83 full-length OPV genomes [7] confirmed the polyphyletic character of the species CPXV and the existence of at least four different clades had been assumed.

**Figure 1:**
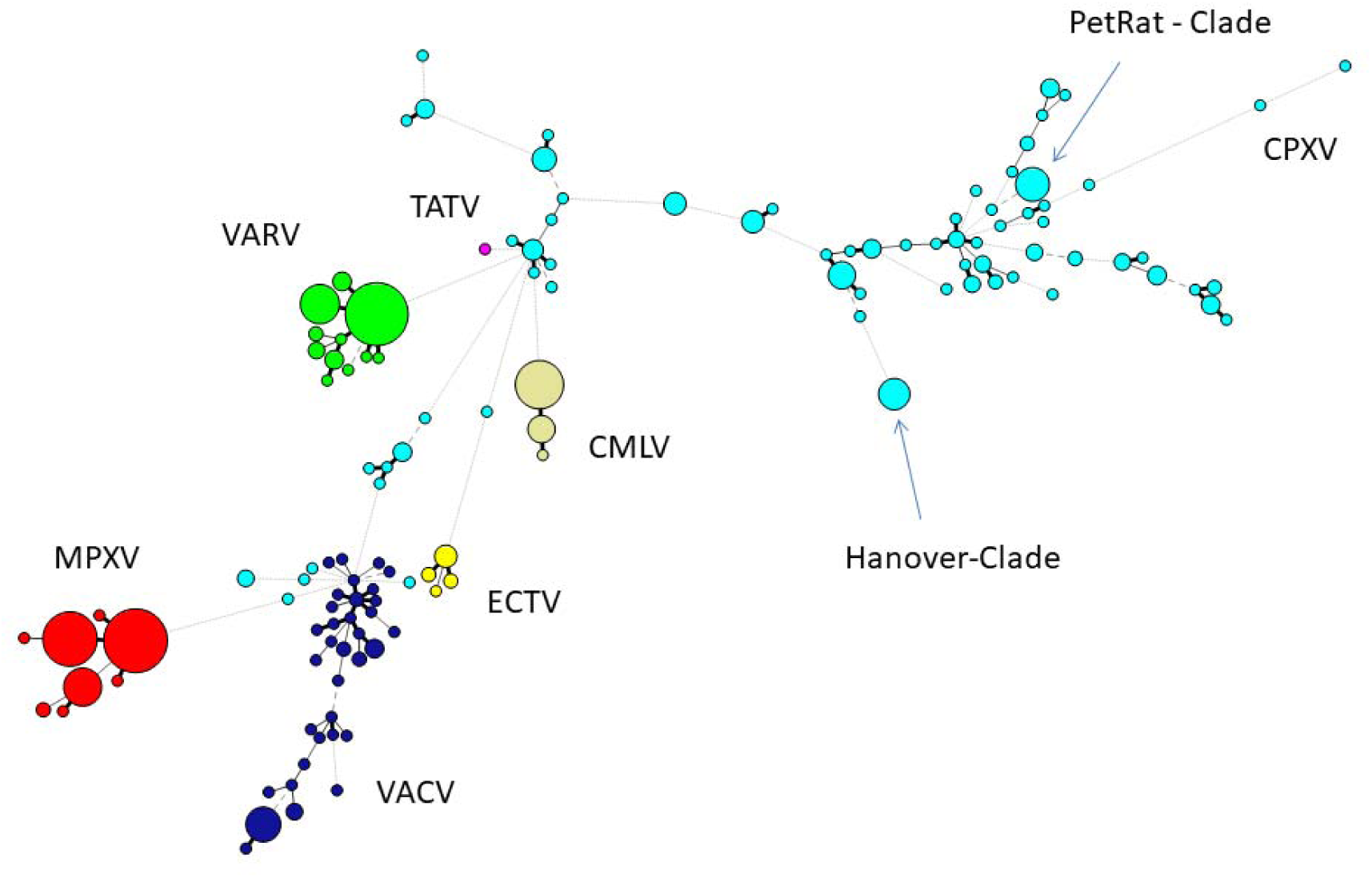
Minimum-Spanning-Tree of 462 hemagglutinin gene sequences of “Old-World”-orthopoxviruses retrieved from NCBI (March 2018). Each circle corresponds to a unique sequence with circle sizes relative to the number of identical sequences. Individual species of the genus Orthopoxvirus are labelled by different colors. CPXV = Cowpox virus, MPXV = Monkeypox virus, VARV = Variola virus, VACV = Vaccinia virus, TATV = Tatera poxvirus, ECTV = Ectromelia virus, CMLV = Camelpox virus.

### Next Generation Sequencing of cowpoxviruses

For each isolate or sample described in this study, one single sequence of at least 200.000 nucleotides with an average sequencing depth of 74 (minimum of 45-fold) could be determined. Each of the sequences spans the entire highly conserved core genome. The outermost terminal inverted repeats [24] could not be reliable determined using a short-read-sequencing technology in combination with the described conservative approach, and were therefore discarded.

In order to determine the reliability of the sequencing method used here on the Illumina platform we re-sequenced two CPXV strains for comparison with their already published genome sequences in order to estimate inter-study reliability. CPXV EP4 Hannover (Table 1) from our in-house collection was compared to Germany_1980_EP4 (HQ420895) which originates from the same isolate, but was sequenced by a different institution using Illumina Sequencing technology as well [25]. The second CPXV strain was Ratpox09 (Table 1, Case #4), whose sequence (LN864565) had already been determined on the Illumina platform by Hoffmann et al. [26] and in this study, too. Not a single nucleotide difference between the sequences could be observed.

In a second approach we wanted to determine whether the process of virus isolation in cell culture would result in nucleotide differences as compared to a sequence directly generated from DNA of lesion material. To this end we sequenced both, two virus isolates obtained from the cats Leo and Sammy (Table 1) and DNA directly extracted from paraffin-embedded lesion material of the same cats. Comparison of the genome of CPXV_2015_5 and DV400026 PF (Leo) as well as CPXV_2015_4a and CPXV_DV400033 PF (Sammy) revealed identical sequences in both cases.

In order to compare the genomes described in this study with already published CPXV genomes, we selected prototypic members of three genomic CPXV-clades described by Franke et al.[7]. Strain HumGra07 (KC813510) belongs to the VARV-like CPXV clade marked in red, strain Finland2000Hum (HQ420893) belongs to the VACV-like CPXV clade marked in green, all other strains EP4, K779, K780, MKY2002, CatBer07, HumLan08 and RatHei09 (HQ420895, AY902281, KY549147, HQ420898, KC813502, KC813492 and KC813504, resp.) belong to the genetic diverse CPXV-1 lineage marked in blue (Figure 2). Whereas HumLan08 and RatHei09 are epidemiologically linked to each other and show identical genome sequences, two genomes from isolates from the city Leipzig in East Germany (2015-1 and 2015-6) [7] [27] originate from two epidemiological unlinked human cases. This was done in order to achieve an idea of the dimension of genetic distances of unrelated samples with the same geospatial and temporal, as well as host attribution. These latter two sequences differ in 111 nucleotide positions from each other.

Strains belonging to HA-sequence type C030 (S1_Table1) are all clustered to the VARV-like strains and strains of HA-sequence type C029 (S1_Table1) show strong similarity to the EP4-strain (HQ420895) within the CPXV-1 lineage.

**Figure 2:**
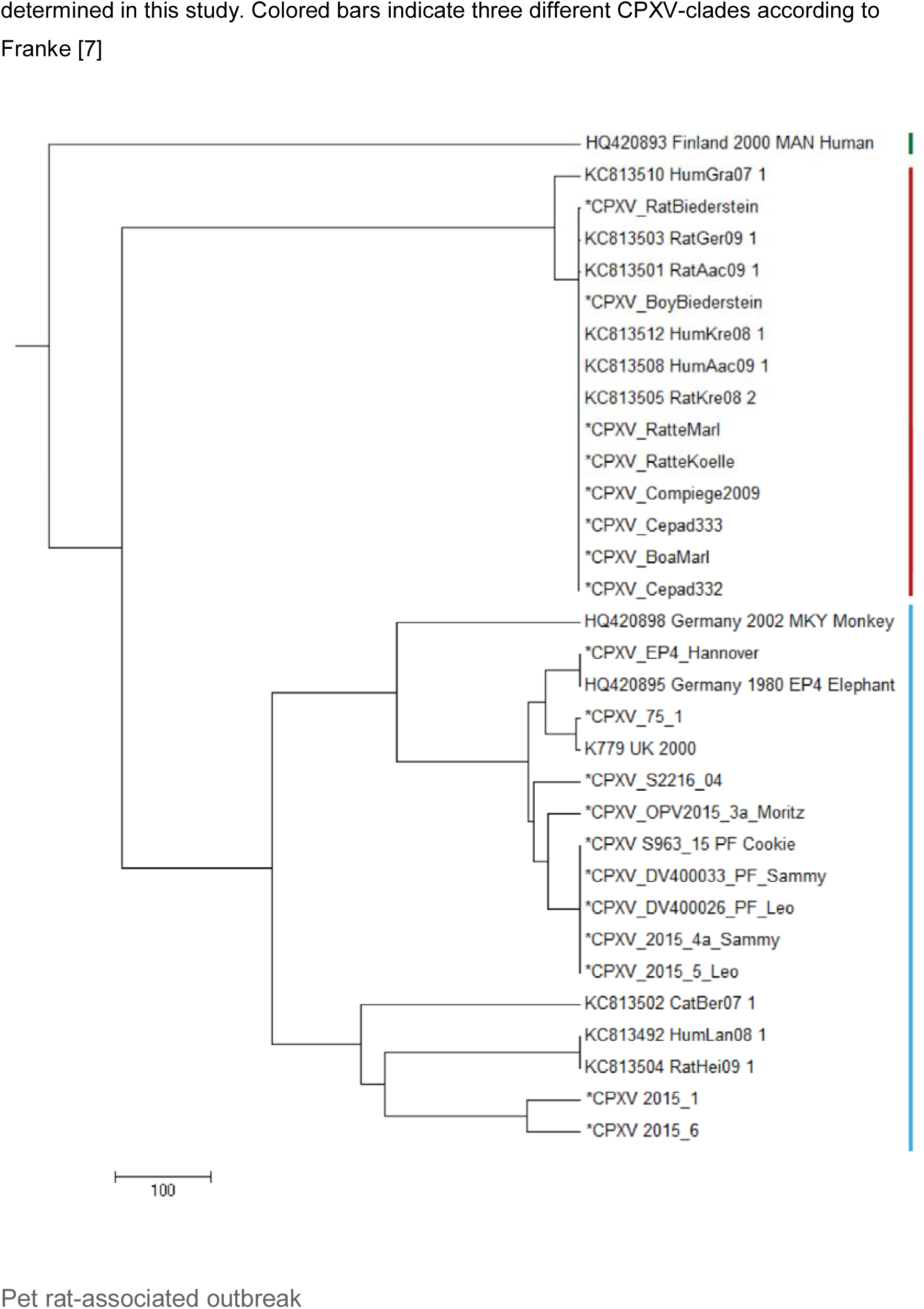
Maximum Parsimony-Dendrogram of extracted and concatenated SNP-positions. Branch lengths show nucleotide differences between the investigated strains. All nodes show bootstrap values of 100%. Sequences marked with an asterisk (*) had been determined in this study. Colored bars indicate three different CPXV-clades according to Franke [7]

### Comparison of sequences generated in this study

In this study we successfully isolated 4 CPXVs from specimens of a boy, his pet rat as well as from a boa and a rat. By including additional four strains of our strain collections, a total of eight pet rat-associated CPXVs genomic sequences were determined. Following alignment a total of 197.964 nucleotide positions could be used for further investigations. Comparison of the eight sequences determined here revealed that six sequences were identical: the strains “Cepad 332” and “Cepad 333” (Case #7) and “Compiegne” (Case #5) had been isolated in France, strains “Boa Marl” and “Rat Marl” had been isolated in the same town in West-Germany (Case #4) and strain “Rat Koelle” originates from South Germany (Case #2). Compared to these six sequences, two sequences ‐ obtained from a sick rat with a proven transmission to a boy (Case #3, South-Germany) ‐ differed in as little as either two or three SNPs, respectively. In gene CPXV089 (according to the nomenclature of the CPXV reference strain Brighton) a nucleotide exchange was leading to an amino-acid change (A → G) and in gene CPXV139 a non-synonymous mutation (R → W) was observed. In addition, strain “Boy Biederstein” has one more nucleotide exchange downstream the gene CPXV035 in an intergenic region. As the young boy was indeed infected by his rat (“Rat Biederstein”), this third and unique mutation of the isolate “Boy Biederstein” indicates a mutually new introduced point mutation during the period of infection.

### Comparison with sequences derived from GenBank

In a next step, we compared our dataset with five sequences already deposited in GenBank from other pet rat-associated CPXVs [8]. A maximum difference of up to eight nucleotides was initially observed between the isolates “HumAac09” and “Rat Biederstein”. These sequences had been generated [8] by using the Roche 454 technology which is known to produce sequencing errors especially in homopolymer regions [28]. Visual inspection and comparison of the alignment confirmed that most differences could be attributed to homopolymer regions. As no raw reads are publically available, a re-calculation with actually available error-correction algorithms could not be performed. By ignoring those positions, strains HumKre08 and RatKre08 from a rat and its owner (Case #1) were identical to the six sequences determined here as well as with strain HumAac09 (Case #3) from a different rat owner. The associated rat isolate “RatAac09” however, differed in two nucleotides as compared to the human isolate. The genetic difference of strain “RatGer09” (Case # 2) was determined to be of two nucleotides. By comparing this pet rat associated-dataset with sequences already published at NCBI, the closest next neighbor was determined to be strain HumGra07 (KC813510), which originates from a case of human cowpox in Graz, Austria, 2007, and differed in a total of 51 SNPs (Figure 3).

**Figure 3:**
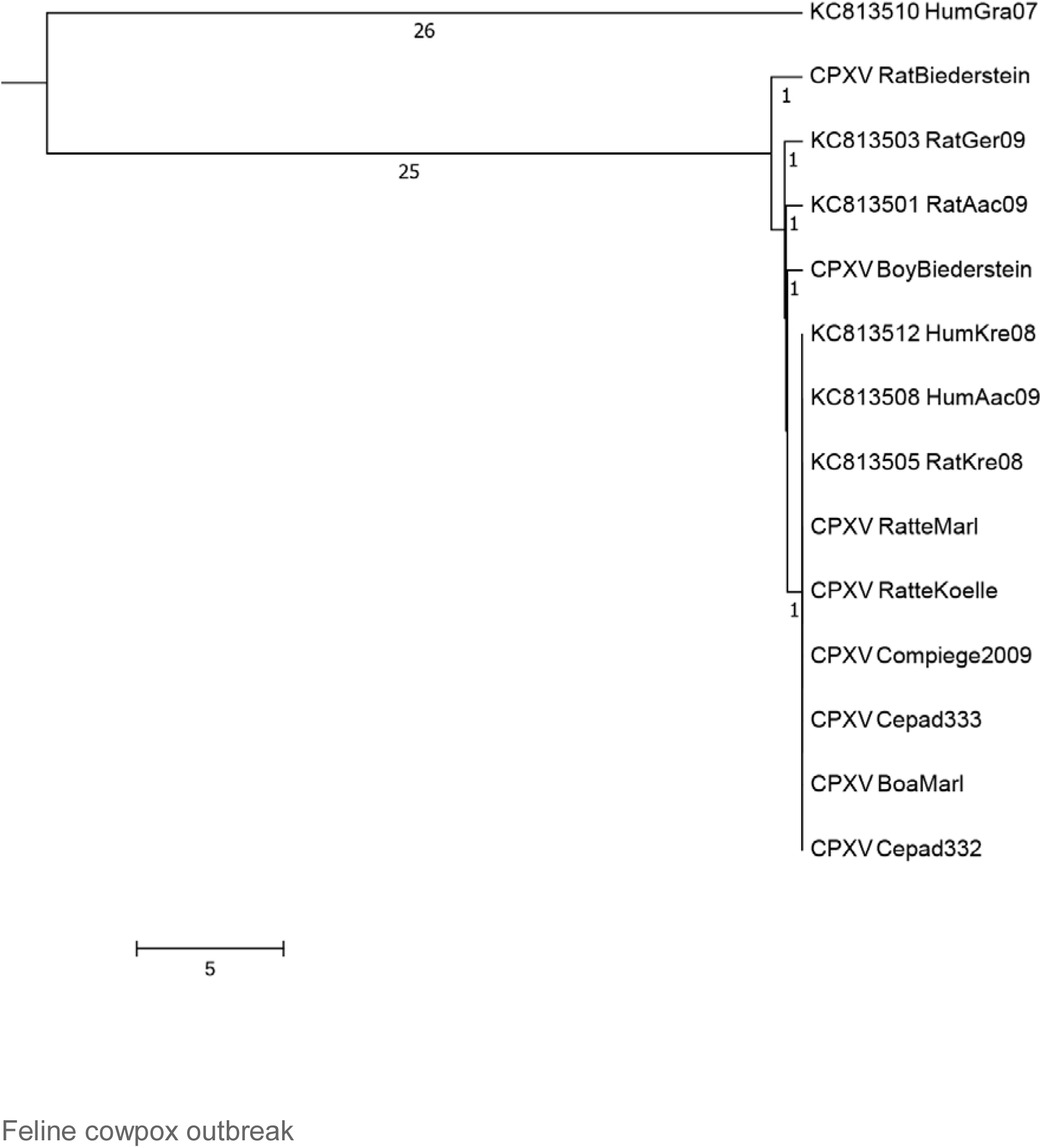
Maximum-Parsimony Dendrogram of extracted and concatenated SNP-positions of pet rat-associated cowpox virus strains. Branch lengths show nucleotide differences between the investigated strains. All nodes show bootstrap values of 100%.

### Comparison of sequences generated in this study

Plaque-like skin lesions had been observed in five cats which had been hospitalized in a period of 5 weeks in a veterinary clinic. CPXV infections were diagnosed and based on identical HA-sequences in four out of five cats (Leo, Sammy, Cookie, and Moritz), it was assumed that infections could have been acquired in the clinic due to contact [15]. DNA extracted from three virus isolates and from three paraffin-embedded tissues (labelled “PF” in figure 4) was used to generate whole genomic sequences. In depth-analysis of the assembly of genomes derived from the paraffin-embedded samples showed a higher initial miscalling rate of nucleotides within reads, similar to observations made before [29]. By using a sequence depth >120-fold, the consensus bases were all called correctly as compared to the sequence derived from isolates. Comparison of all genomes demonstrated that five genomes were identical. The sixth sequence derived from cat Moritz (CPXV_OPV2015_3a) differed in 65 positions (Figure 4) with SNPs are located throughout the whole genome.

### Comparison with sequences derived from GenBank

In order to find a possible relationship to previously reported cases of cowpox, sequences generated here were compared to genomes with identical HA sequences. A total of 201.073 nucleotide positions could be used for evaluation. Strain “CPXV S2216/04” which was isolated from a cat in 2004 in the same region, differed in 96 SNPs and strains CPXV “K779 UK2000” and “CPXV 75/1” differed in 108 nucleotides.

**Figure 4:**
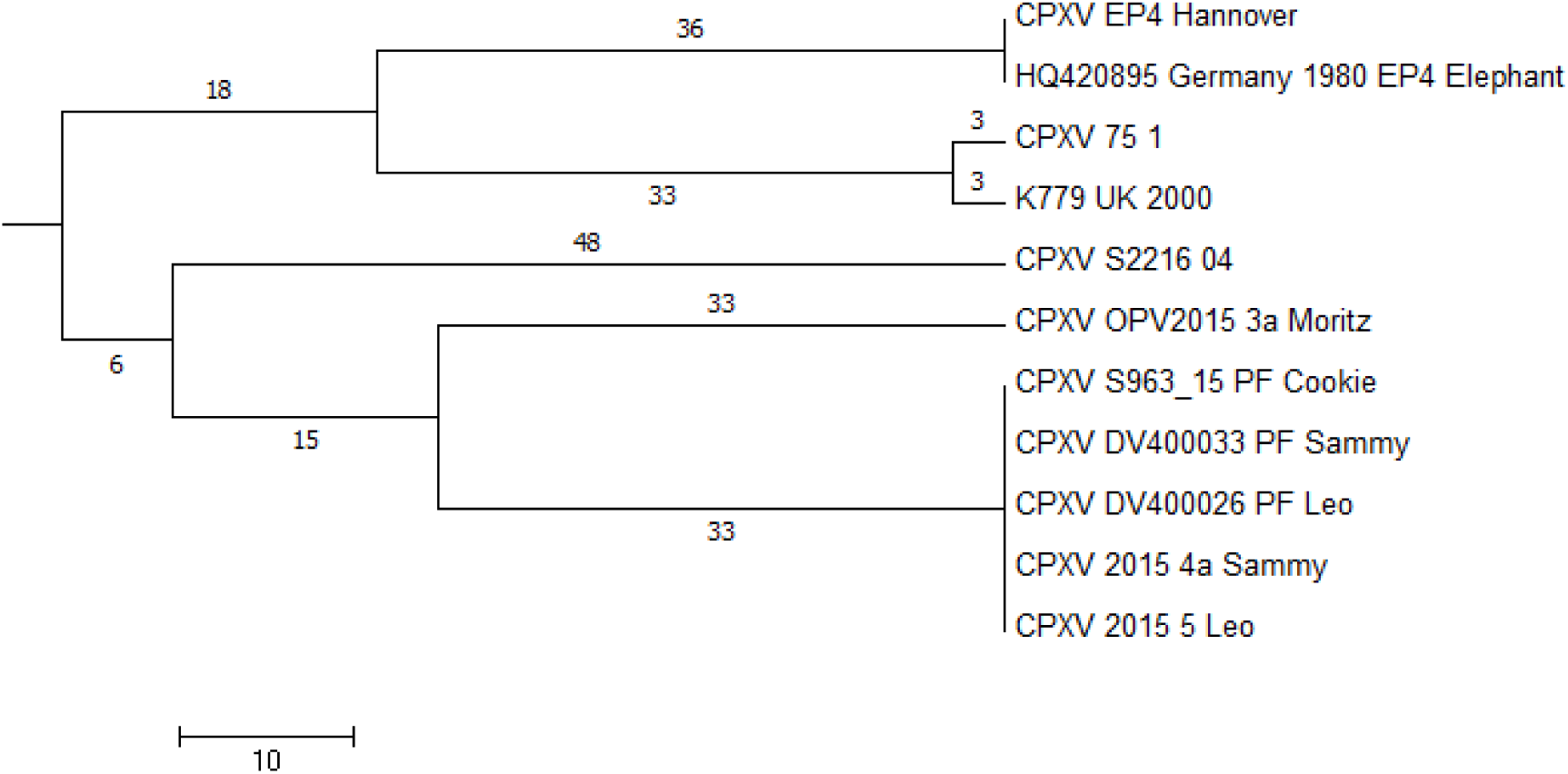
Maximum-Parsimony Dendrogram of extracted and concatenated SNP-positions of feline cowpox-associated strains. Branch lengths show nucleotide differences between the investigated strains. All nodes show bootstrap values of 100%.

## Discussion

Conducting PCR screening assays targeting the HA gene in combination with sequencing of the HA ORF allow (i) unambiguous identification of orthopoxviruses and (ii) assigning a given isolate to an OPV species. Sequencing can be performed even in laboratories, which do not have access to a routine NGS-pipeline but rely on Sanger sequencing. Here, we aligned 447 HA sequences of orthopoxviruses and could show that the HA genetic signatures of VARV, MPXV, CMLV and ECTV are indeed species-specific. This can be used as a useful tool for rapid typing of new isolates and assigning them to a given OPV species without conducting time-consuming and laborious phenotypic characterizations. Typing of and differentiating the species CPXV and VACV is also possible with the restriction that some CPXV strains might be mistyped as VACVs (Figure 1 and Table S1, CPXV groups C001-3, C053 and C063). According to whole genome analysis, these strains form a separate clade referred to as “VACV-like” CPXV [7] which has a common ancestor with the species VACV. According to the species definition of the International Committee on Taxonomy of Viruses the taxonomic classification of CPXVs has to be revisited to avoid confusion. Current data support the recognition of five monophyletic clades of cowpox viruses as valid species [6].

The discriminatory power of the HA gene sequence is excellent despite it is a short sequence of about 940 nucleotides: HA analysis allows to unambiguously identify VARV, MPXV, ECTV, CMLV, VACV and CPXV. In case of VARV and MPXV further differentiation is possible: variola major strains can be differentiated from variola minor strains and Western African MPXVs can be differentiated from Central African MPXVs.

In contrast, the heterogeneity of CPXVs is much higher with 79 unique sequences out of 144 strains (Figure 1 and Figure S1). The existence of unique isolates can be best explained by evolution over time in a local rodent reservoir and strains therefore cluster according to their geographic origin. This is a prerequisite for conducting phylogeographic attributions and trace back analyses to understand cowpox outbreaks. Researchers have taken advantage of the sequence diversity to investigate cowpox outbreaks and conducted trace-back analysis. For example, in a rather small outbreak involving a rat, an elephant and the animal caretaker three identical HA sequences were obtained. Since the sequence had not been reported before, a rat-elephant-human transmission was assumed [9]. In another outbreak with generalized infections in banded mongooses (*Mungos mungo*), jaguarundis (*Herpailurus yagouaroundi*) and a time-delayed case of human cowpox, an identical and unique HA sequence indicates a common source of infection, i.e. infected food rats [13]. If HA sequences are distinct, then it can be concluded that they are not epidemiologically linked, if HA sequences are identical, they could or could not be epidemiologically linked. This scenario merits to be further investigated using a supplementary method [9], [13]. To this end we have investigated two HA sequence groups (C029 and C030; Table S1) which are composed of 11 and 14 identical sequences, respectively, by determining the sequence of the entire genome which has the highest discriminatory power.

Sequence group C030 consists of various specimens derived from seven pet rat-associated cases (Table 1). We could show here, that all 14 genomes had maintained an extremely high level of clonality. The same high level of clonality (maximum of 3 SNPs) was observed in 4 genomes obtained from a smallpox outbreak, which spread form in Iran in 1968 to Syria and Yugoslavia in 1972. A highly efficient proofreading mechanism of the viral DNA-polymerase and/or a low selective pressure in the respective hosts might be the reason. The origin of the pet rat outbreak could not be identified with certainty: close to the German-Czech border a die-off, especially of young rats was observed in 2009 at a rat breeder facility and the outbreak in Munich (case #2) could be traced back to this facility [14]. The authors [11] claim that the young rats in France (case #7) in 2011 had been imported from the Czech Republic. Further attempts involving Czech authorities to unveil the origin of feeder or pet rats and their trading routes failed.

Sequence group C029 consists of a total of 11 identical HA sequences (Table 2), four sequences from an outbreak of feline cowpox and seven sequences from single cases of cowpox which had occurred in the larger area of Hannover and in the UK in previous years [8], [25], [30]. The latter cases are not linked to the feline cowpox outbreak. Genome sequencing clearly proved that a hospital-acquired transmission had occurred in three out of four cats. In combination with epidemiological data [15], cat Leo could be identified as the index case with few nodular skin lesions at initial presentation. At this time an infection with CPXV was not considered and Leo was hospitalized. During this time Sammy with a history of a car accident and Cookie, which was treated for renal insufficiency, were hospitalized as well. A fourth cat, Moritz (with an identical HA sequence) was also hospitalized. All cats could have been potentially exposed. However, the genome of Moritz differed in 61 nucleotides compared to identical genomes of Leo, Sammy and Cookie. We conclude that Moritz has not become infected in the clinic.

Here, we demonstrated that traceback analyses based on HA sequences only are not sufficient to detect the true chains of transmission in an outbreak. Whole genome sequencing is the method of choice since it provides the highest discriminatory power and in combination with epidemiological data outbreak investigations can be performed.

For orthopoxviruses the specific viral load within lesions is fortunately very high, so that even with smaller sequencing instruments, sufficient sequencing depth can be achieved using clinical samples. Even sequencing the entire genome of FFPE treated lesion material can be performed.

With use of Next-Generation-Sequencing, knowledge of genetic epidemiology of orthopoxviruses has increased during the last two to three years [6], [7], [26]. Creating and updating reliable databases of orthopoxvirus genomes would even allow a better understanding of genetic diversity and clonality. In combination with new sequencing technologies, this knowledge might help to be prepared for future orthopoxvirus outbreaks of MPXV or CPXV. This has already successfully applied during the recent Ebola outbreak in West Africa: Sequences were generated on site [31] [32], linked to clinical cases and enabled outbreak reconnaissance and containment of the outbreak.

## Corresponding Author

Markus Antwerpen,

## Data Availability

All sequence data is available at NCBI under BioProject PRJNA369073

## Acknowledgment

**Funding**

**This publication was supported by the European Virus Archive goes Global (EVAg) project that has received funding from the European Union’s Horizon 2020 research and innovation programme under grant agreement No 653316.**

**This study was also supported in part by the European Union’s Horizon 2020 research and innovation program under grant agreement No 643476 (COMPARE).**

